# Hypoxia-induced Complement Component 3 Promotes Aggressive Tumor Growth in the Glioblastoma Microenvironment

**DOI:** 10.1101/2024.01.28.577617

**Authors:** Rebecca Rosberg, Karolina I. Smolag, Jonas Sjölund, Elinn Johansson, Christina Bergelin, Julia Wahldén, Vasiliki Pantazopoulou, Crister Ceberg, Kristian Pietras, Anna M. Blom, Alexander Pietras

**Affiliations:** Division of Translational Cancer Research, Department of Laboratory Medicine, Lund University, Lund, Sweden; Section of Medical Protein Chemistry, Department of Translational Medicine, Lund University, Malmö, Sweden; Division of Medical Radiation Physics, Department of Clinical Sciences, Lund University, Lund, Sweden

## Abstract

Glioblastoma (GBM) is the most aggressive form of glioma with a high rate of relapse despite intensive treatment. Tumor recurrence is tightly linked to radio-resistance, which in turn is associated with hypoxia. Here, we discovered a strong link between hypoxia and local complement signaling using publicly available bulk, single cell, and spatially resolved transcriptomic data from human GBM patients. Complement component 3 (*C3*) and the receptor *C3AR1* were both associated with aggressive disease and shorter survival in human glioma. In a genetically engineered mouse model of GBM, we found C3 specifically in hypoxic tumor areas. In vitro, we found an oxygen level-dependent increase in *C3* and *C3AR1* expression in response to hypoxia in several GBM and stromal cell types. Presence of C3 increased proliferation of GBM cells under hypoxic conditions, as well as clonal survival of GBM cells following radiation. Targeting C3aR using the antagonist SB290157 decreased GBM cell self-renewal in vitro, and prolonged survival of glioma bearing mice both alone and in combination with radiotherapy while reducing the number of M2-polarized macrophages. Our findings establish a strong link between hypoxia and complement pathways in GBM, and support a role of hypoxia-induced C3a-C3aR signaling as a contributor to glioma aggressiveness.

## Introduction

Glioblastoma (GBM) is the most common and aggressive primary brain tumor in adults [1]. Despite intensive treatment with surgery, irradiation, and chemotherapy, all patients suffer recurrence of treatment-resistant tumors [2, 3]. The overall survival for patients remains low, with a median survival of less than 2 years [4]. GBM is characterized by abundant pseudopalisading necroses, that are surrounded by areas of severe hypoxia [5]. Tumor hypoxia is associated with tumor angiogenesis and a metabolic shift towards anaerobic glycolysis, as well as tumor cell stemness and radioresistance [6–8]. Although the malignant cell response to hypoxia is well characterized, heterotypic cell-cell signaling involving stromal cells in the hypoxic niche remains poorly understood.

Prominent non-malignant cell types including microglia/macrophages [9], astrocytes [10], and endothelial cells [11], play tumor-supportive roles in the GBM microenvironment. Within the tumor, stress related to e.g. hypoxia, chemotherapy, and radiotherapy is not restricted to the tumor cell compartment, but is likely to affect the phenotype and behavior of the entire spectrum of cell types in the microenvironment. Indeed, we have previously shown that radiotherapy results in a net tumor-supportive microenvironment in the brain [12, 13]. Specifically, tumor-associated astrocytes respond to radiotherapy [12] and severe hypoxia [14] with a reactive phenotype, that in turn promotes aggressive tumor growth. Similarly, radiotherapy results in an altered immune environment in part because of radiation-induced phenotypic shifts of microglia and macrophages [15]. It is likely that hypoxia-induced gene expression changes result in altered cell-cell communication between stromal and tumor cells.

Complement component 3 (C3) is a key marker of astrocyte reactivity [16–18], and a central component of the complement system of innate immunity, found at high levels in serum. Most circulating complement proteins from serum are restricted from entering the brain due to the blood brain barrier (BBB), thus most of complement proteins in the brain are expressed locally or enter through a leaky BBB [19, 20]. When activated by proteolysis, C3 cleavage products, including C3a, can signal via various receptors, affecting cellular activation, inflammation, and metabolism [21]. However, its presence and function in tumor-associated gliosis remains undetermined. C3a-C3aR signaling has been implicated in brain development, neural plasticity, and protection against hypoxic-ischemic brain injuries [22–24], suggesting the potential of a role also in the setting of brain tumors. Here, we used publicly available transcriptomic data, a genetically engineered mouse model of GBM, and primary cultures of GBM and stromal cell types to investigate the presence and function of C3 in the GBM microenvironment, and its role in relation to hypoxia.

## Materials and Methods

### Sex as a biological variable

Animal, cell line, and clinical samples representing both sexes were used.

### Generation of Murine Gliomas

Gliomas were induced in Nestin/tv-a (Ntv-a) mice by intracranially injecting RCAS-PDGFB- and RCAS-shp53-transfected chicken fibroblast DF-1 cells (ATCC ® CRL-12203™, ATCC, Manassas, VA, USA) in the neonatal brain, as previously described [25]. Mice were monitored daily and sacrificed upon development of glioma symptoms. For C3aR-antagonist studies, mice were randomized into groups and treated daily (5 days on, 2 days off) with 1mg/kg SB290157 trifluoroacetate (Tocris, 6860/5) intra-peritoneally, starting at the occurrence of glioma symptoms (day 37 +/-1) for up to a total of 30 injections. Radiotherapy was performed after three days of injections of SB290157 at day 40 (+/-1). Mice were sedated using isofluorane, and cranial radiotherapy was administered with a 10 mm field in one single fraction of 10 Gy on a 220 kV preclinical research platform (XenX, XStrahl Inc, Suwanee, GA, USA).

### Cell Culture and Treatments

Primary human astrocytes (3H Biomedical, Uppsala, Sweden) were cultured in astrocyte medium (3H Biomedical) supplemented with 2% FBS and 1% PenStrep solution. U3082MG, U3065MG and U3084MG glioma cells were obtained from HGCC (hgcc.se) and were cultured in HGC medium on laminin coated plates as described previously [26]. Cells were passaged using Accutase (Thermo Fisher Scientific, Waltham, MA, USA). Glioma cell line U251MG (Sigma) and HMC3 (ATCC) microglia were cultured in DMEM (Corning) supplemented with 10% FBS (Biological Industries) and 1% PenStrep (Corning). Cells were cultured at 37°C in a humidified incubator containing 21% O2 and 5% CO2, unless otherwise stated. Hypoxia was generated in a Whitney H35 Hypoxiastation (Don Whitley Scientific) or an InvivO2 400 Hypoxia Workstation (Baker Ruskinn, Bridgend, United Kingdom).

### Proliferation assay

Cells were seeded at a density of 1x104 cells per well in a 96-well plate, incubated with 180 ng/ml serum purified C3. C3 was purified from human EDTA plasma using two consecutive anion exchange columns Q-Sepharose and DE52 cellulose. C3 was 95% pure as judged by gel electrophoresis. Plates were incubated either at 21% or 0.1% O2 as previously described. Cell viability/cell growth were measured at 24, 48 and 72 h by addition of WST-1 (Abcam, ab155902) to the medium. Plates were read at 450 nm in a Syngery 2 Plate reader (BioTek).

### Colony formation assay

U251MG cells were plated at 150 cells/well in a 6-well plate. At day 2, cells were irradiated with 4 Gy using a CellRad x-ray irradiator (Faxitron), after 1 hour with or without addition of 180 ng/mL human serum purified C3. Cells were then allowed to incubate until visible colonies were formed, or up to 14 days. Colonies were fixed in 4% PFA, followed by Crystal Violet 0.01% staining for 1 h, and imaged in LAS-3000 system. Colonies were counted using Image J/Fiji (version 2.1.0).

### Immunofluorescence

Human glioma cells (U3082MG) were seeded on cover slip glass coated with poly-orthinine and laminin and allowed to attach. Cells were rinsed 3 times with phosphate-buffered saline (PBS) followed by addition of fresh medium with or without 180 ng/ml human serum purified C3 and were placed at 0.1% oxygen for 72 h. Cells were fixed in 4% paraformaldehyde (PFA) for 20 minutes followed by washing and permeabilization using 0.3% Triton X-100 (Sigma-Aldrich) in PBS. Cells were blocked using 2.5% fish gelatin in a solution of 0.05% Tween20 in PBS followed by incubation with primary anti-Ki67 (ThermoFisher, rat, 14-5698-82), incubated over-night at 4°C. Cells were incubated with the appropriate secondary antibodies in presence of Hoechst 33342 (Sigma-Aldrich) for 1 h. Olympus BX63 microscope with DP80 camera and cell-Sens Dimension v 1.12 software (Olympus Corporation, Tokyou, Japan) were used to capture images. Ki67+ cells were quantified for each cell using CellProfiler 4.0.6 [27].

### Multiplexed immunofluorescence

Animals were euthanized upon glioma symptoms, and whole brains were embedded in Optimal cutting temperature compound (OCT) (Thermo Fisher Scientific), and frozen in ice-cold isopentane. Sections were air-dried for 30 min followed by fixation in ice-cold acetone. Permeabilization was performed in 0.3% Triton X-100 in PBS. Sections were blocked in 1% BSA followed by primary antibody incubation.

To examine the C3 expression we designed a custom multiplexed immunofluorescence detection panel using Alexa Fluors and antibodies targeting C3 (Hycult, HM1065, 1:150 dilution), Olig2 (R&D Systems, AF2418, 1:200 dilution) as a tumor marker, GFAP for detection of astrocytes (Abcam, ab4674, 1:400 dilution) and Nestin (Abcam, ab7659, 1:100 dilution) for detection of perivascular niches. After PBS wash, the following species-specific, fluorophore-conjugated secondary antibodies were used to reveal antibody staining (FITC, Alexa Fluor 594, Alexa Fluor 647 and Alexa Fluor 750, 1:500 dilutions). Cell nuclei were labelled with 4′,6-diamidino-2-phenylindole (DAPI) (Sigma-Aldrich). To control for nonspecific signals, tissue sections were incubated with secondary antibodies alone. All antibodies were optimized for concentrations using monostainings imaged on an Olympus BX63 microscope. Scanning of the full antibody panel was performed on PhenoImager (Akoya Biosciences). A spectral fluorophore for each primary antibody and autofluorescence library was made to enable optimal multispectral unmixing. Spectral unmixing of images was performed using the inform software package (Akoya Biosciences).

Staining of brains from SB290157-treated mice was performed as described above, with Olig2, GFAP, Ki67 (Thermofisher, RM9601-S) and the microglia/macrophage marker F4/80 (BioRAD, MCA497) combined with the following secondary antibodies (FITC, Alexa Fluor 555, Alexa Fluor 594, Alexa Fluor 647, 1:500). For vessel detection, staining was performed using CD34 (Thermofisher, 14-0341-81) and for M2-macrophages using CD206 (Invitrogen, MA5-16868).

### Real-Time qPCR

RNA was isolated using the RNeasy Mini Kit together with the Qiashredder Kit (QIAGEN) according to the manufacturer’s protocol. cDNA was synthesized using random primers and Multiverse transcriptase enzyme (Applied Biosystems). Amplifications were run using a QuantStudio 7 real-time PCR system (Applied Biosystems) with SYBR Green Master Mix (Applied Biosystems). Melting curves were run for each primer pair to ensure specificity of primers. Relative gene expression was normalized to the expression of three housekeeping genes (*SDHA*, *UBC*, and *YWHAZ*) using the comparative Ct method [28]. All detections were performed in triplicates.

### Sphere Forming Assays

For primary sphere formation, U3082MG cells were dissociated with Accutase then plated at single-cell density 150 cells/well in a 6-well plate in 2.5mL of HGC media. The spheres were grown until visible spheres were formed (up to 14 days) with control or treatments with 50 and 250nM C3aR antagonist (Santa Cruz Biotechnology, SB290157). For the Extreme Limiting Dilution Assay (ELDA), primary U3082MG spheres were dissociated, then replated at 150-900 cells/mL in 300 μL in eight wells of a 48-well dish followed by serial 1:2 dilution across eight columns in the presence or absence of treatment until visible spheres were formed. Any well containing a sphere was scored as positive. Data combined from four biological repeats were quantified using the ELDA analysis tool [29].

### Patient cohort analysis

Data from the Allen Institute for Brain Science IVY-GAP (http://glioblastoma.alleninstitute.org) [30] and the TCGA (TCGA_GBM and TCGA_LGG [31]) were analyzed using the Gliovis data portal [32]. Pan-TCGA data were analyzed using the Xena platform [33]. The results published in here are in whole or part based upon data generated by the TCGA Research Network: https://www.cancer.gov/tcga. Molecular signatures for HALLMARK_HYPOXIA (M5891) [34] and HALLMARK_COMPLEMENT (M5921) [34] in TCGA data were analyzed using GEPIA 2 (http://gepia2.cancer-pku.cn/#dataset) [34].

### 10x Visium spatial transcriptomics analysis

Previously published human GBM spatial transcriptomic data sets [35], available at https://datadryad.org/stash/dataset/doi:10.5061/dryad.h70rxwdmj) were processed according to the standardized pipeline as provided by Seurat [36] (https://satijalab.org/seurat/articles/spatial_vignette.html) in R. In short, data were normalized using SCTransform and principal component analysis and UMAP dimensionality reduction were all done using default parameters. We used the R package msigdbr ([37], version 7.5.1) to obtain the hypoxia and complement Hallmark gene sets and and calculated average z-scores of the genes in each specific Hallmark signature in the spots of the GBM spatial transcriptomic Visium data. The Pearson correlation values (p values were corrected for multiple hypothesis testing using the Holm method) between the hypoxia and complement gene set expression scores in each spot were calculated using the R package correlation ([38], version 0.8.4). The hypoxia and complement z-scores were visualized onto the Visium tissue sections using Seurat’s SpatialFeaturePlot function and the results from correlation analyses were visualized using the R package ggpubr ([39], version 0.6.0)

### Single cell RNA sequencing analysis

The processed single cell RNA sequencing dataset comprising 110 primary human GBMs [40] was downloaded from cellxgene (https://cellxgene.cziscience.com/collections/999f2a15-3d7e-440b-96ae-2c806799c08c). Dimensionality reduction coordinates and cell type annotations were as provided by the authors. All data were imported and analyzed using Seurat. 2D UMAPs of clusters, gene expression levels, and hypoxia and complement AddModuleScores (a function in Seurat to calculate gene set module scores) were visualized using the scCustomize package ([41], version 0.7.0) and the SCpubr package ([42], version 2.0.2). Seurat’s FindMarkers function (logfc.threshold = -Inf, min.pct = -Inf, min.diff.pct = -Inf) was used to obtain differentially expressed genes between the positive (expression > 0.0) and negative (expression <= 0.0) cell populations. For the gene set enrichment analysis (GSEA), all genes were ranked by their estimated log2 fold changes. The analysis, as implemented in the fgsea package ([43], version 1.29.1), was performed using the Hallmark gene sets with 5% FDR and 1000 permutations and visualized using ggplot2 ([44], version 3.4.4).

### Statistical analysis

Unless otherwise stated, all values represent mean ± SEM for at least three independent experiments. All statistical analysis were performed in GraphPad Prism vs. 9.3.1. or R-Studio vs 2023.09.0+463. Statistical tests and number of replicates are indicated in Figure legends. Three levels of significance were used.

### Study approval

All animal procedures were conducted in accordance with the European Union directive about animal rights and procedures were approved by Lund ethical committee (M-16123/19).

### Data availability

The data and materials that support the findings of this study are available from the corresponding author upon reasonable request.

## Results

### C3 is associated with aggressiveness in glioma

Bulk RNA-sequencing from TCGA datasets on GBM and low-grade glioma (TCGA_GBM and TCGA_GBMLGG) showed that *C3* expression increased with glioma grade (Fig. 1A), IDH-wt status (Fig. 1B), and was expressed at higher levels in GBM compared to non-tumor brain (Fig. 1C). *C3* expression correlated with worse survival in patients with glioma (Fig. 1D), as well as in IDH-wildtype GBM (Fig. 1E). We generated murine GBM using the RCAS/tv-a system to express *PDGFB* and shRNA targeting *Tp53* in Nestin-expressing cells of the neonatal Nestin/tv-a mouse brain. Tumors were stained for C3, GFAP as an astrocytic marker, and Olig2 for the bulk of tumor cells. In line with TCGA data demonstrating higher levels of C3 in GBM as compared to non-tumor brain tissue (Fig. 1C), we found the vast majority of C3 present within the tumor core, with virtually no detectable stain in the tumor-adjacent healthy brain tissue (Fig. 1F).

**Figure 1.**
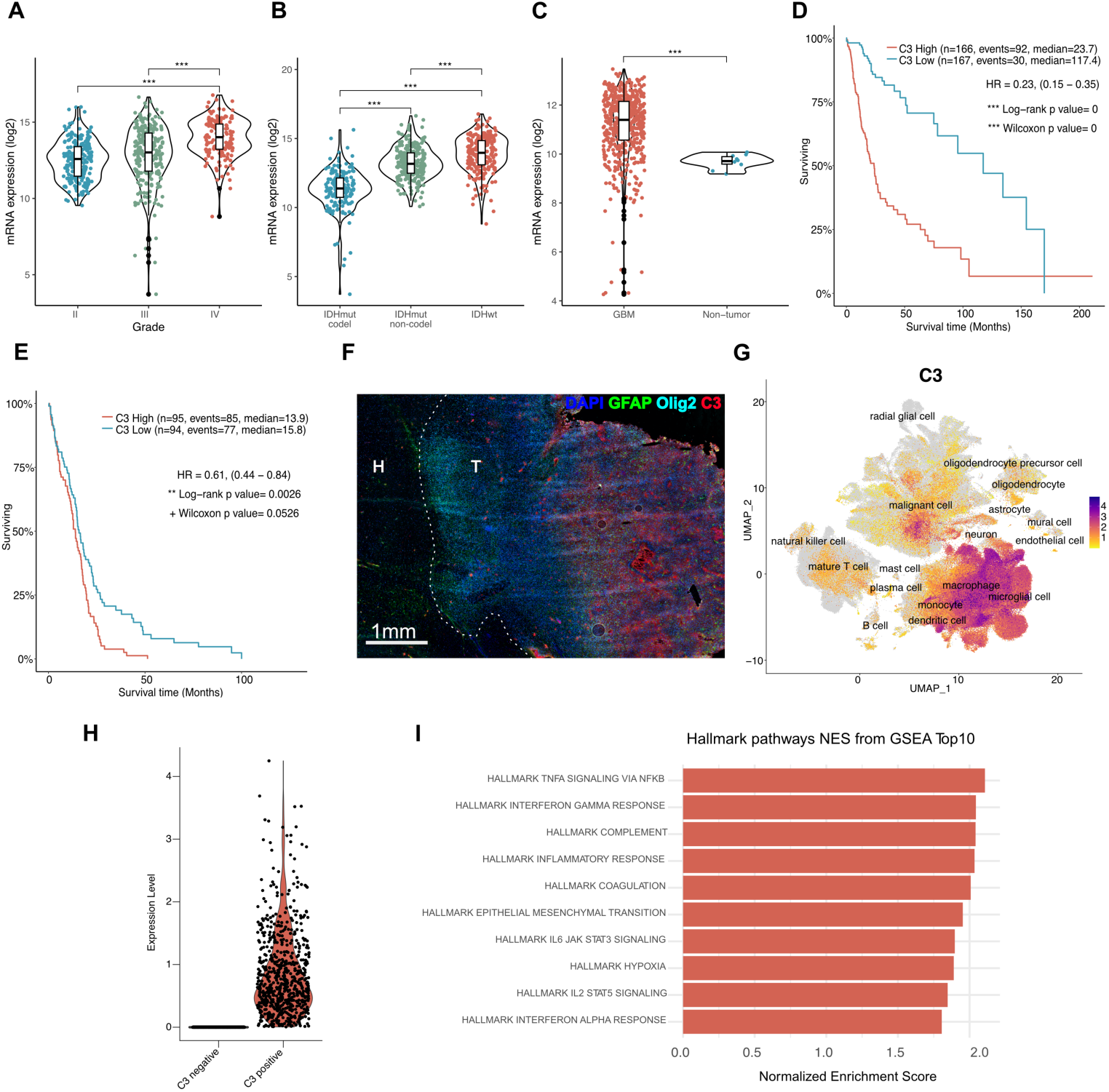
C3 is associated with aggressive GBM. **A**. TCGA data analyzed for C3 expression in glioma grades II, III and IV. **B.** TCGA data analyzed for C3 expression in IDH wildtype (IDHwt) glioma compared to IDH mutant (IDHmut) with or without 1p/19q codeletion. **C.** TCGA data analyzed for C3 expression in GBM compared to non-tumor. **D.** Kaplan-Meier curve showing survival of glioma patients with either high (red) or low (blue) C3 expression based on TCGA data. **E**. Kaplan-Meier curve showing survival of IDH-wildtype GBM with high (red) or low (blue) C3 expression based on TCGA data. **F**. Murine GBM stained for C3, Olig2, GFAP, and DAPI as indicated. T indicates tumor and H indicates healthy brain tissue. **G.** UMAP displaying C3 expression in single cell sequencing data from 16 independent datasets compiled in GBmap [40]. **H.** Malignant cell population divided into C3-low or -high-expressing cells. **I.** Gene set enrichment signature pathways associated with C3 expression in malignant cells. Red colored bars indicate significant Benjamini-Hochberg adjusted P values (padj < 0.05). *, P < 0.05, **, P < 0.01, or ***, P< 0,001. Statistical tests were one-way ANOVA, or unpaired T-test (in case of comparison between two groups) with Tukey post-hoc test.

We next queried the highly integrated GBmap from Ruiz-Moreno et al. [40], comprising 16 independent single cell RNA sequencing data sets, and a total of over 330,000 cells from 110 GBM patients (Supp. Fig. 1A)[40]. Analysis showed that *C3* is expressed by several cell types, including macrophages, microglia, astrocytes, endothelial, T-cells, and tumor cells (Fig. 1G). We classified cells within each cell type into either *C3*-negative or *C3*-expressing cells (Fig. 1H, Supp. Fig. 1B-D), and asked whether certain gene sets were associated with *C3* expression. Gene set enrichment analysis revealed that *C3*-expressing cells in all tested cell types (Astrocytes, (Supp. Fig. 1B, Supp. Fig. 1E), Mural Cells, (Supp. Fig. 1C, Supp. Fig. 1F) and Macrophages/Microglia (Supp. Fig. 1D, Supp. Fig. 1G)) showed an enrichment for gene signatures associated with inflammatory processes such as TNF-alpha and IFN-γ-signaling (Fig. 1I). Interestingly, *C3*-expressing malignant cells were associated with gene sets for EMT and hypoxia (Fig. 1I). Based on the entire dataset, cells that scored positive for *C3* expression showed increased expression of several genes involved in activation of the classical complement pathway (*C1QA, C1QB, C1QC* and *C3*), as well as Factor D (*CFD*) from the alternative complement pathway (Table 2). Early complement inhibitory genes *CD46*, *CD55* and *CSMD1* were downregulated (Table 2). Interestingly, late complement inhibitors such as *CD59* were also increased in *C3*-expressing cells, suggesting that *C3*-expressing cells may still be protected from cellular lysis mediated through the membrane attack complex (MAC), while there is a potential for early complement activation to take place (Table 2).

**Table 1.**
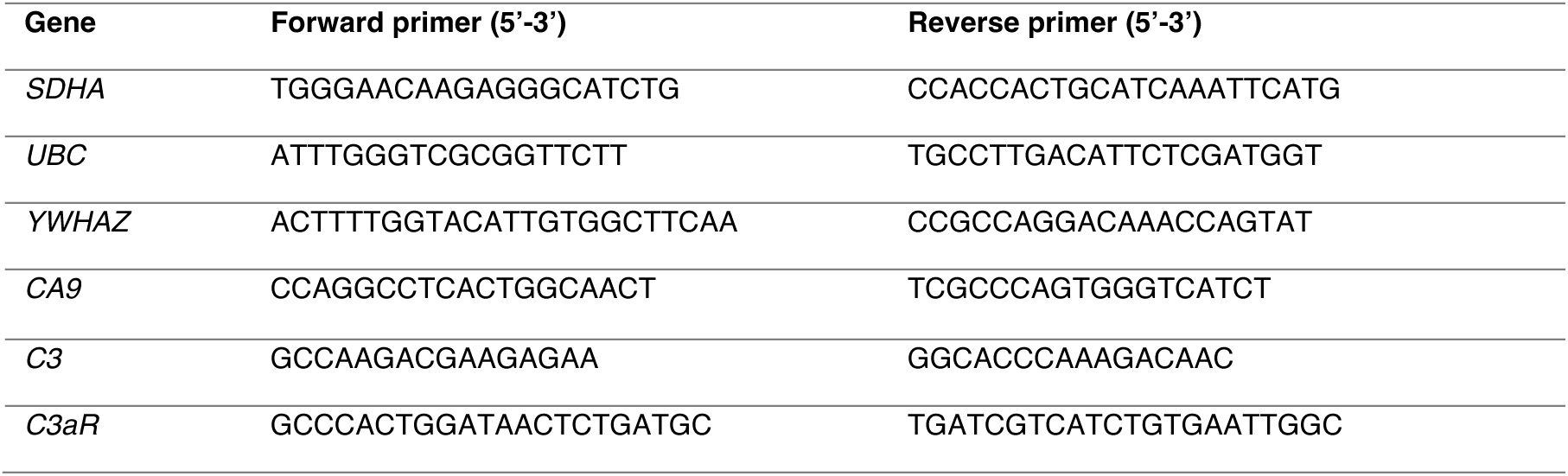
Primers sequences for RT-qPCR.

**Table 2.**
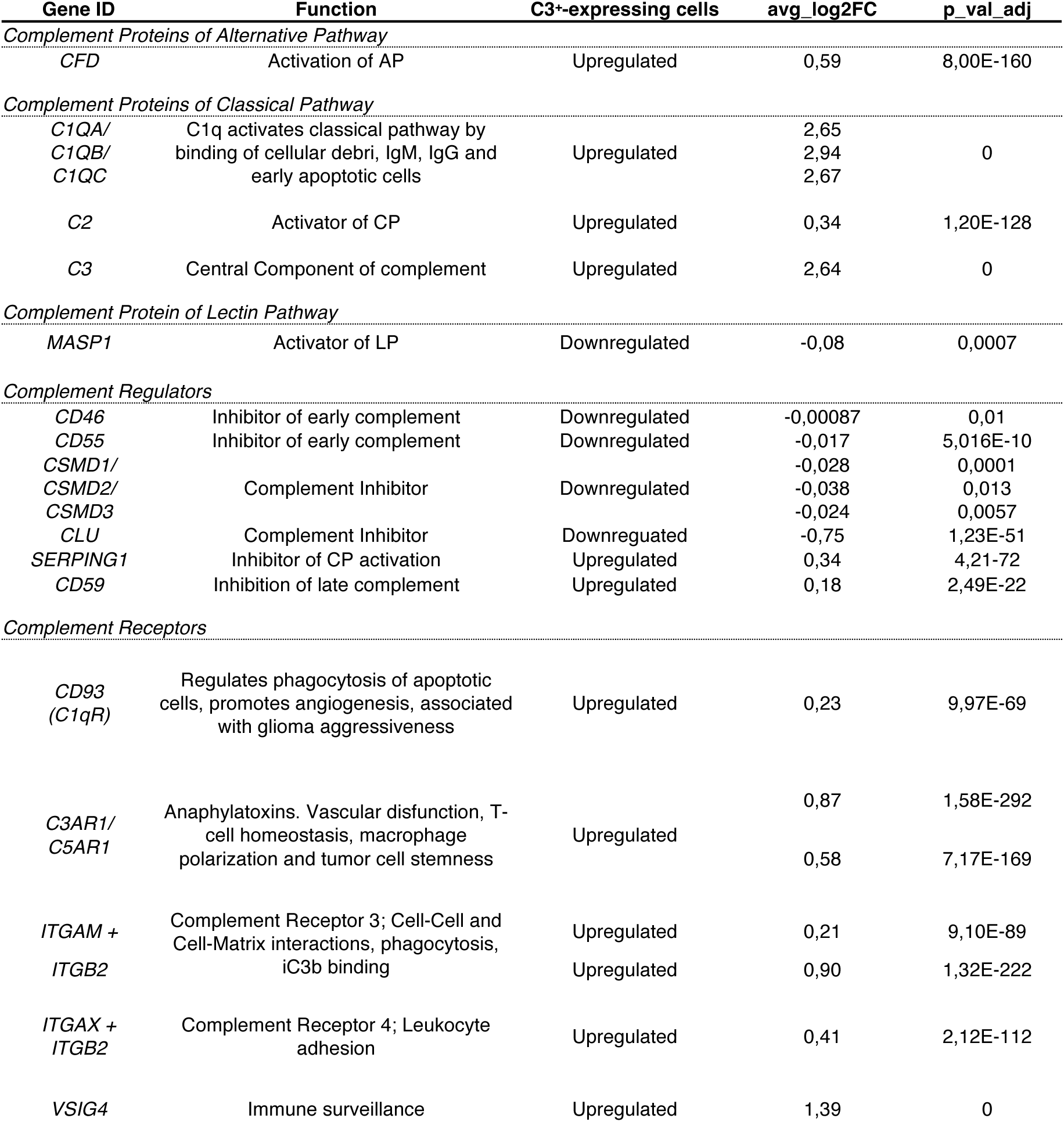
Gene expression of complement proteins in C3-expressing cells.

### Hypoxia is associated with local complement signaling in GBM tumors

To understand the role of C3 and complement in the tumor microenvironment, we returned to stain the murine GBMs (Fig. 1F) for C3, GFAP, Olig2, and Nestin to highlight perivascular tumor areas. C3 was present primarily in perivascular (Fig. 2A) and perinecrotic (hypoxic) (Fig. 2B) tumor niches, in close proximity to Nestin-expressing cells. Considering the high C3 levels found in hypoxic tumor areas and the association between C3-expressing GBM cells and the hypoxia gene signature (Fig. 1I), to further investigate the relationship of complement expression to hypoxia, we applied the Hallmark_complement and Hallmark_hypoxia gene expression signatures to a publicly available dataset of spatial transcriptomics from 19 human GBM patients [35]. These analyses revealed a strong spatial correlation of the two signatures (Fig. 2C-F), with the complement signature almost exclusively present in tumor areas scoring high for the hypoxia signature. Bulk RNA sequencing from TCGA confirmed a strong correlation of the signatures in both low-(Fig. 2G) and high-grade gliomas in larger patient cohorts (Fig. 2H). In the human GBM single cell dataset (GBmap), we found that the hypoxia (Supp. Fig. 1B) and complement (Supp. Fig. 1C) signatures are primarily expressed by cells derived from the myeloid linage, such as macrophages, monocytes, microglial cells, as well as endothelial cells, astrocytes and to some extent tumor cells themselves in human GBM (Supp. Fig. 2A-D). We next sought to determine whether *C3* is directly upregulated in response to hypoxia. Astrocytes, microglia, and human primary GBM cells, were cultured in various oxygen tensions (normoxia (21% O2), hypoxia (1% O2) and severe hypoxia (0.1% O2)) for 72 h. Hypoxia induction was confirmed by increased levels of *CA9* (Fig. 2I-K). Several of the stromal cell types tested (Fig. 2I-J), and GBM cell lines (Fig. 2K) showed increased *C3* expression, as well as increased expression of its receptor, *C3AR1*, under hypoxic conditions. Together, these data show that local C3 expression in GBM is highly associated with hypoxia, and that growth under hypoxic conditions itself can cause upregulation of *C3* and in most instances also the *C3AR1* receptor.

**Figure 2.**
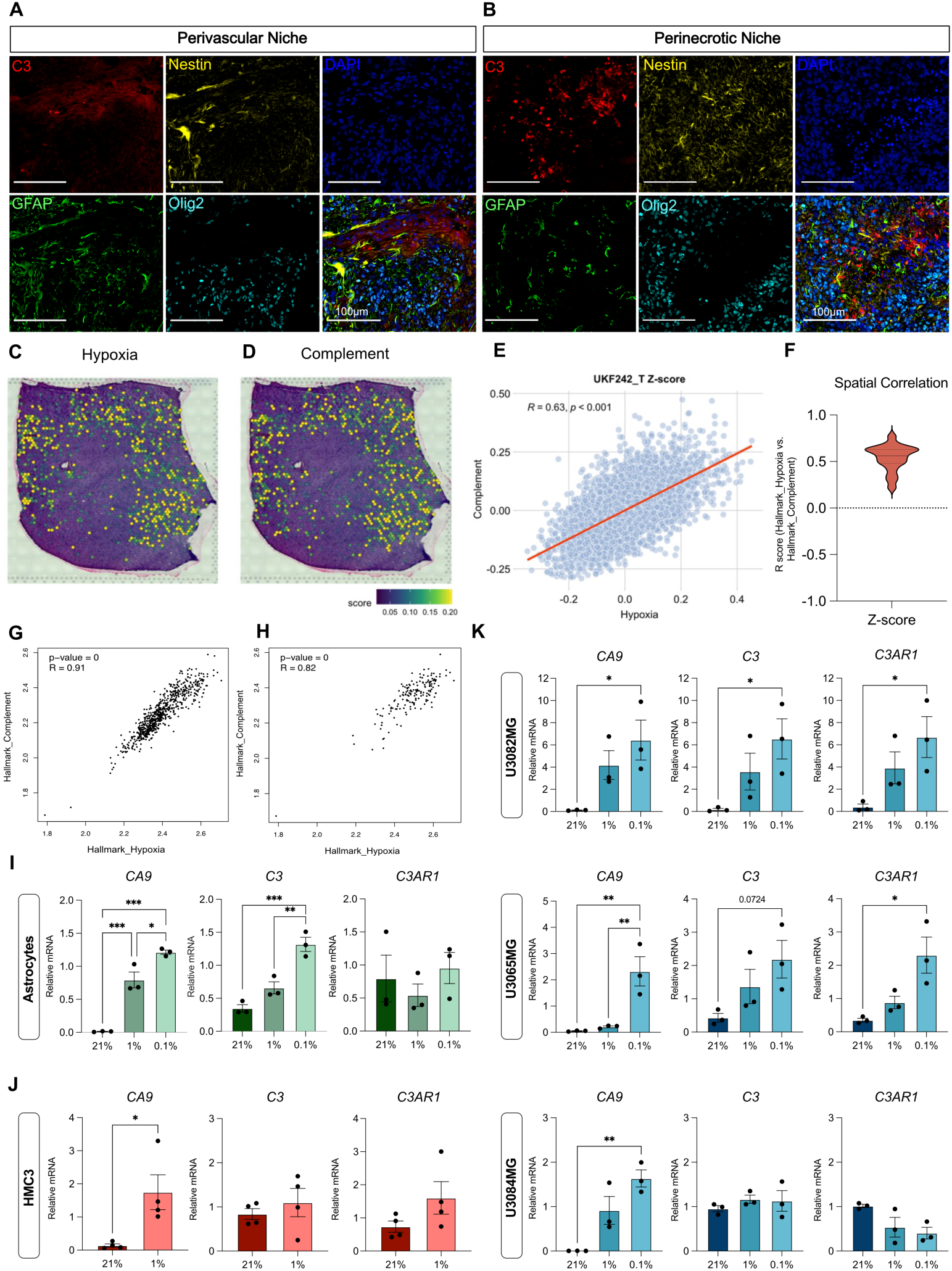
Hypoxia is associated with local complement signaling in GBM tumors. **A-B.** Immunofluorescent detection of C3 (red), Nestin (yellow), Nuclei (DAPI, blue), GFAP (green) or Olig2 (cyane) in the perivascular (A) or hypoxic (B) niches of murine RCAS-PDGFB- and RCAS-shp53-induced gliomas. **C-D.** Hallmark_Hypoxia (C) and Hallmark_Complement (D) signatures mapped on spatially resolved transcriptomics from human GBM samples, displayed with Z-score. **E.** Spatial correlation between Hallmark_Hypoxia and Hallmark_Complement in one representative tumor (UKF242). P values were corrected for multiple hypothesis testing using the Holm method. **F.** Distribution of R-values for the correlation between Hallmark_Hypoxia and Hallmark_Complement in a total of 19 human GBM tissue samples with an average correlation score of R=0.54 [35] **G-H.** Pearson correlation coefficient between Hallmark_Complement and Hallmark_Hypoxia signatures in the TCGA GBMLGG (G) or TCGA GBM (H) dataset. **I-K.** Expression of CA9, C3, and C3AR1 mRNA in human primary astrocytes (n=3), HMC3 microglia (n=4), or primary human glioma cells U3082MG (n=3), U3065MG (n=3) or U3084MG (n=3) in response to normoxia (21% O2), hypoxia (1% O2) or severe hypoxia (0.1% O2). Error bars represent SEM. *, P < 0.05, **, P < 0.01, or ***, P< 0,001. Statistical tests were one-way ANOVA, with Tukey post-hoc test, or unpaired t-test in case of two sample comparisons.

### C3 promotes proliferation and survival of glioma cells

To determine whether presence of C3 can modulate tumor cell properties, we added serum-purified human C3 in serum-free culture medium of U3082MG primary glioma cells. Proliferation (Fig. 3A) as well as the proportion of Ki67+ cells (Fig. 3B-C) increased in presence of C3 specifically under hypoxic conditions, when *C3AR1* was upregulated (Fig. 2K). In a clonal survival assay following a single dose of 4 Gy irradiation, U251MG glioma cells formed significantly more colonies when C3 was present in the culture medium (D-E). Together, these data further support a role for C3 as a pro-survival factor during cellular stress such as hypoxia and irradiation [45, 46].

**Figure 3.**
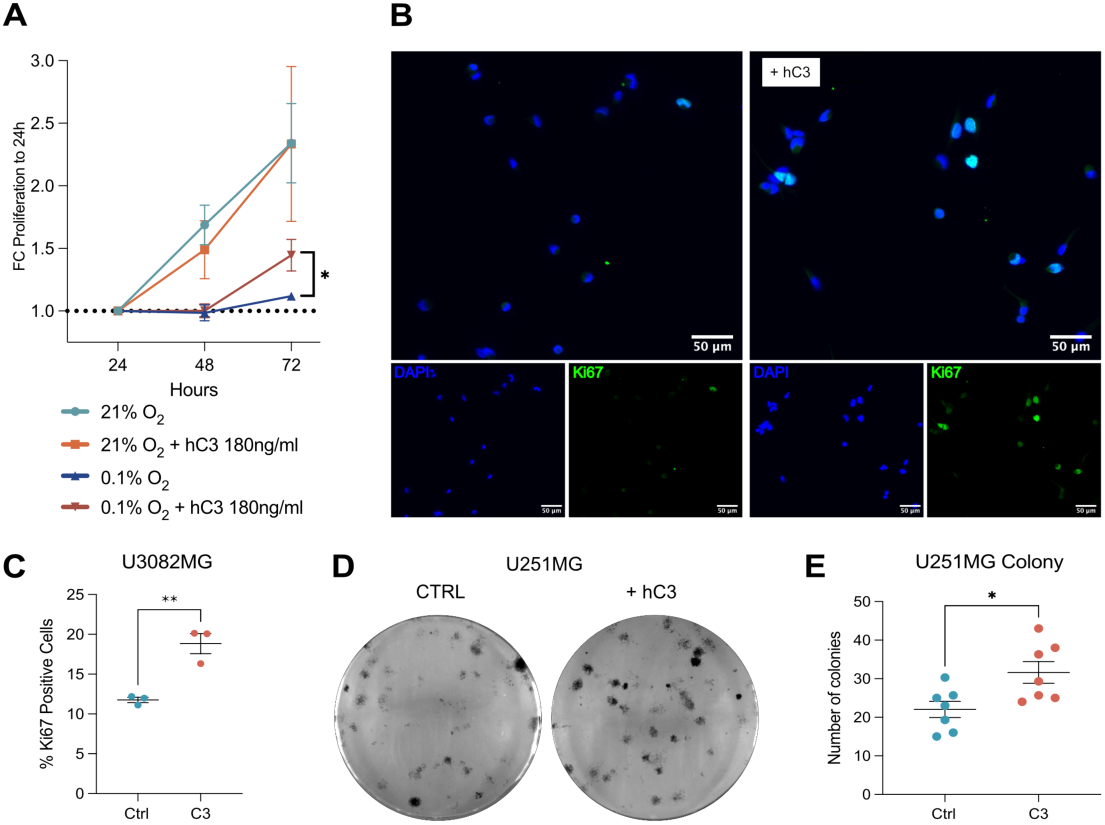
C3 supports glioma cell growth under stressful conditions. **A.** U3082MG glioma cell proliferation under normoxic (21% O2) and severe hypoxic (0.1% O2) conditions with or without presence of human serum-purified C3 (n=5) as measured by the WST-1 assay. **B.** Representative images of immunofluorescent detection of Ki67 positive cells in U3082MG glioma cells grown under severe hypoxia (0.1% O2) with or without presence of C3 (180 ng/ml). **C.** Quantification of Ki67+ U3082MG cells (n=3). **D.** Representative image of clonal survival of U251MG glioma cells after 4 Gy irradiation with or without presence of human C3 (180 ng/ml) in the medium in triplicate wells. **E.** Quantification of number of colonies (n=7). Error bars represent SEM. *, P < 0.05, **, P < 0.01 from unpaired t-tests.

### *C3AR1* is highly expressed in GBM and is associated with aggressive tumor growth

We next sought to investigate the expression of complement receptors in GBM using TCGA data. We found that *C3AR1* expression is higher in GBM as compared to all other cancers analyzed in TCGA (Fig. 4A), whereas Complement Receptor 1 (*CR1*), Complement Receptor 2 (*CR2*), C1qR (*CD93*), and *C5AR1* were expressed at varying levels (Supp. Fig. 3A-D). *C1qR* (CD93) (Supp. Fig. 3E), *C3AR1* (Figure 4B), and *C5AR1* (Supp. Fig. 3F), but not *CR1* (Supp. Fig. 3G) or *CR2* (Supp Fig. 3H), were highly upregulated in GBM compared to normal brain tissue. *C3AR1* increased with glioma grade (Fig. 4B) and IDH wildtype status (Fig. 4C), and was expressed at higher levels in GBM as compared to non-tumor brain (Fig. 4D). Furthermore, *C3AR1* was associated with shorter survival in glioma (Fig. 4E) as well as in IDH-wildtype GBM (Fig. 4F). Single cell RNA sequencing data from GBmap showed that *C3AR1* was expressed mainly by cells of the immune compartment, such as microglia/macrophages and mature T-cells (Fig. 4G). Furthermore, the tumor cells that were positive for *C3AR1* scored high for signatures associated with inflammation, as well as several pathways associated with cell cycle regulation (Fig. 4H-I). Because of the high expression of *C3AR1* in GBM, and the pro-tumoral effects of C3 in GBM cell cultures, we tested the effect of the C3aR antagonist SB290157 in the extreme limiting dilution assay -a functional assay measuring the self-renewal capability of U3082MG glioma cells. U3082MG cells treated with SB290157 showed a reduced ability to form spheres (Fig. 4J), suggesting that signaling through C3aR contributes to glioma cell self-renewal.

**Figure 4.**
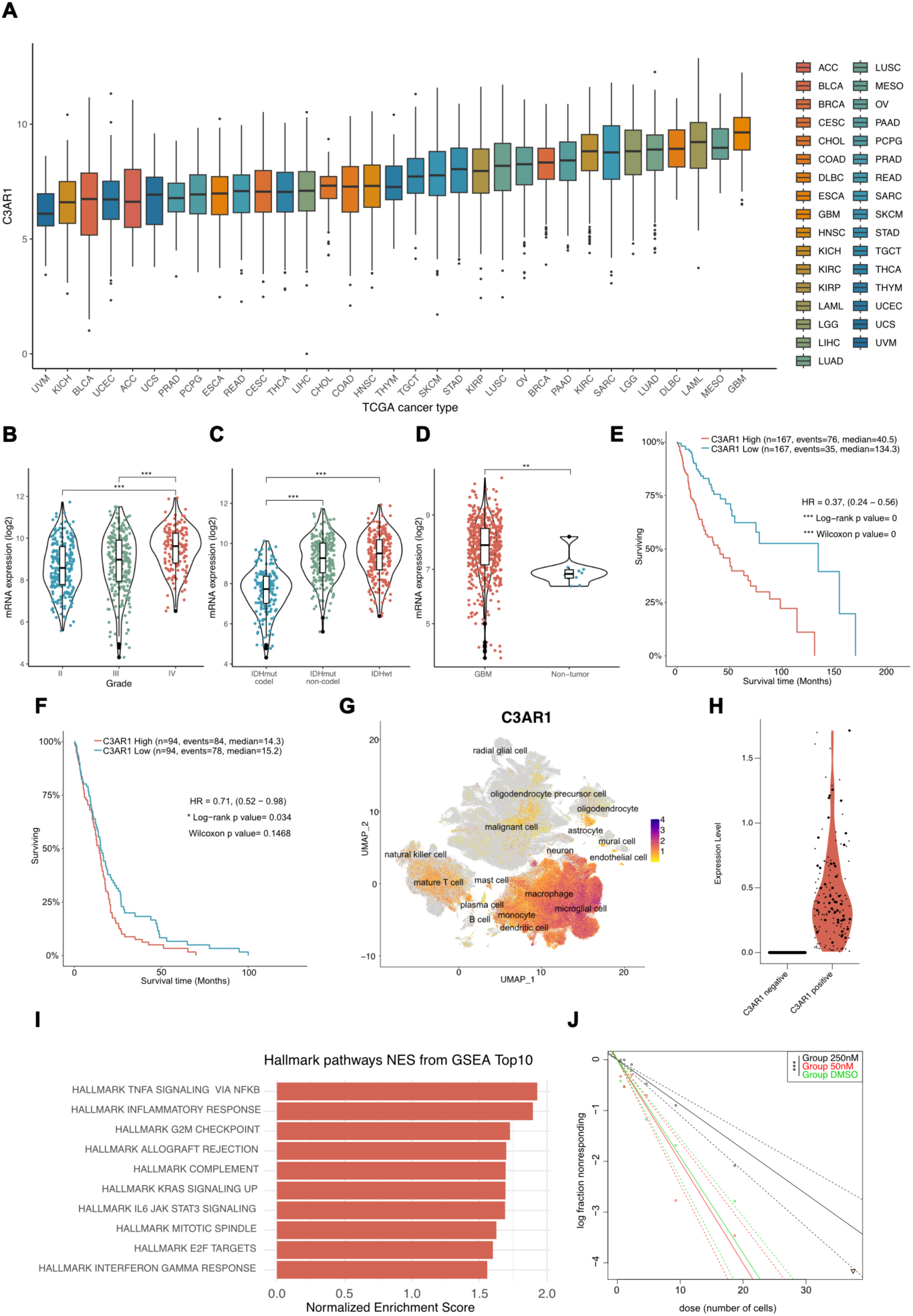
C3AR1 is associated with aggressive GBM. **A.** C3AR1 expression of Pan-Cancer TCGA data of common cancer types (n=33). **B.** C3AR1 expression in relation to glioma grade as analyzed in TCGA data. **C.** C3AR1 expression in IDH wildtype glioma compared to IDH mutant with or without 1p/19q codeletion (Tukey post-hoc test) as analyzed in TCGA data. **D.** C3AR1 expression in GBM compared to non-tumor as analyzed in TCGA data. **E.** Kaplan-Meier curve showing survival of glioma patients with either high (red) or low (blue) C3AR1 expression based on TCGA data. **F**. Kaplan-Meier curve showing survival of IDH-wildtype GBM with high (red) or low (blue) C3AR1 expression based on TCGA data. **G.** UMAP displaying C3AR1 expression in single cell sequencing data from 26 independent datasets compiled in GBmap [40]. **H.** C3AR1 expression of malignant cells divided into C3AR1-expressing or non-expressing cells. **I.** Geneset enrichment analysis of the C3AR1-expressing malignant cells. Red colored bars indicate significant Benjamini-Hochberg adjusted P values (padj < 0.05). **J.** Log-fraction plot of a combination of independent extreme limiting dilution sphere formation assays (n=4) of U3082MG glioma cells treated with SB290157. *, P < 0.05, **, P < 0.01, or ***, P< 0,001. One-way ANOVA, or unpaired T-test (in case of comparison between two groups) with Tukey post-hoc test.

### C3aR is a potential therapeutic target in glioma

To investigate C3a-C3aR-signaling as a possible therapeutic target in GBM, we generated murine tumors as previously described (Fig. 1F). Daily intraperitoneal injections (5 days on, 2 days off) of 1mg/kg of the C3aR-antagonist SB290157 were initiated at the time of emergence of brain tumor symptoms, with or without radiotherapy (10 Gy) (Fig. 5A). We found that mice treated with the C3aR-antagonist alone displayed increased survival (Fig. 5B), similarly to what was observed with radiotherapy alone (Fig. 5B). Furthermore, mice treated with a combination of the C3aR-antagonist and radiotherapy showed further increased survival time as compared to radiotherapy alone (Fig. 5B). Immunostaining (Fig. 5C) of treated tumors showed no difference in the expression of F4/80 (Fig. 5D), Olig2 (Supp. Fig. 4A), GFAP (Supp. Fig. 4B) or Ki67 (Supp. Fig. 4C) in SB290157-treated tumors. Interestingly, tumors treated with SB290157 displayed fewer M2-polarized CD206+ macrophages (Fig. 5E-F), despite the lack of significant differences in the total macrophage/microglia population (Fig. 5D), suggesting that C3a-C3aR1 signaling can modulate macrophage polarization. Most CD206+ cells were located in perivascular tumor areas or in association with blood vessel-like structures (Fig. 5E). Tumors treated with a combination of radiotherapy and SB290157 were less vascular in general, as measured by the amount of CD34+ cells, but no such difference was apparent in non-irradiated tumors (Supp. Fig. 4E-F). Together, these data suggest that C3aR is a potential therapeutic target in the GBM microenvironment, targeting of which might affect the cellular composition of tumors in general, and macrophage polarization states specifically.

**Figure 5.**
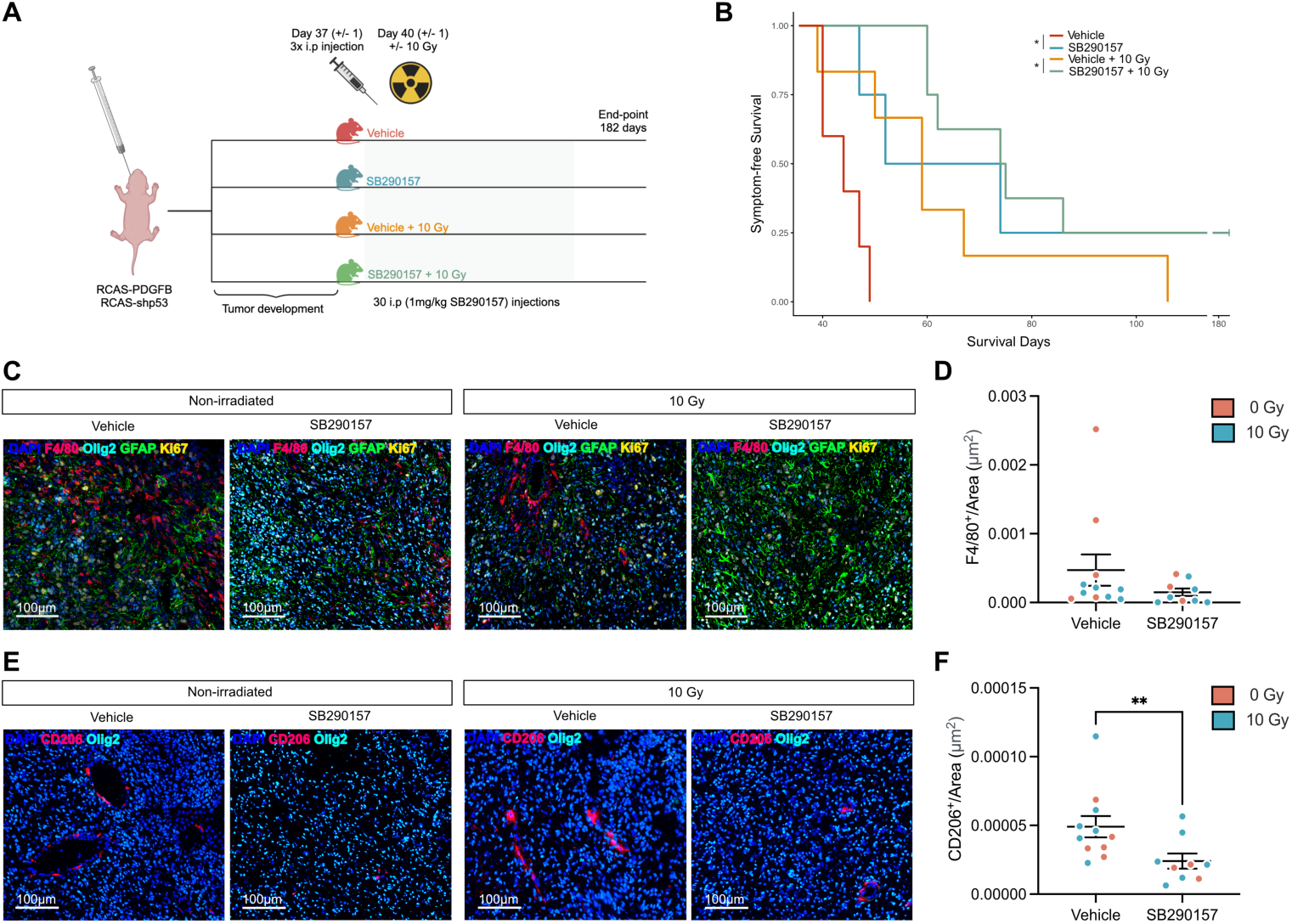
C3AR1 is a possible therapeutic target in GBM. **A.** Illustration (created with BioRender.com) of study design for treatment with SB290157 (1mg/kg) with or without combination with 10 Gy radiotherapy of murine gliomas induced through RCAS-PDGFB and RCAS-shp53 in Nestin/tv-a mice. **B.** Kaplan-Meier curve showing the survival of mice treated with Vehicle (red, n = 5), SB290157 (blue, n = 4), Vehicle + 10 Gy (orange, n = 6) or SB290157 + 10 Gy (green, n = 8) as indicated. **C.** Representative image of immunofluorescent staining of macrophages/microglia (F4/80), tumor cells (Olig2), and astrocytes (GFAP) in mice treated as indicated. **D.** Quantitative analysis of F4/80+ cells/Area (μm2) as indicated. **E.** Representative images of immunofluorescent staining of tumor cells (Olig2) and CD206+ macrophages/microglia in mice treated as indicated. **F.** Quantitative analysis of CD206+ cells/Area (μm2) as indicated. *, P < 0.05, **, P < 0.01, or ***, P < 0,001. Statistical analysis was performed using the Mann-Whitney U-test.

## Discussion

GBM recurrence is driven by treatment-resistant cells, which in turn are popularly believed to be associated with certain tumor niches, such as hypoxic or perivascular niches [47]. Much effort has been directed towards characterizing the tumor cell response to hypoxia, but stromal cell responses to hypoxia and their potential contribution to tumor progression remain relatively unexplored. We have previously shown that astrocytes adopt a reactive phenotype with tumor-promoting properties in response to hypoxia and irradiation. In this study, we show that cells of the GBM microenvironment upregulate the central protein of the complement system, C3, under hypoxic conditions. Presence of C3 during cellular stress, such as hypoxia or irradiation, led to increased survival of glioma cells, supporting previous findings that C3 can act as a pro-survival factor during cellular stress [46, 48, 49].

We demonstrated that complement pathway gene expression is almost exclusive to tumor areas enriched for hypoxia-inducible gene expression in human GBM. While it is likely that this correlation is driven largely by the fact that the majority of complement gene expression in gliomas is derived from tumor-associated macrophages and microglia, cell types known to be enriched in hypoxic tumor areas [50], our experimental work established that at least *C3* and *C3AR1* could be directly induced in numerous cell types when cultured under hypoxic conditions. These findings add another level of complexity to previous reports linking hypoxia with the complement system. Olcina, et al. demonstrated that tumors harboring mutations in the complement pathway displayed elevated hypoxia signaling [51]. Furthermore, several genes associated with the complement system have been shown to be regulated by hypoxia in various cancer cell lines [52, 53].

Complement activation is a hallmark of several cancers. In particular, generation of C3a and C5a fragments has been associated with a tumor- and metastasis-supportive microenvironment. It is thought that malignant cells hijack complement regulatory pathways, resulting in chronic early complement activation, immunosuppression, angiogenesis, proliferation, EMT, and pro-tumorigenic signaling [54, 55], while managing to escape complement-mediated lysis by upregulating late complement inhibitory proteins. Although each cancer can exhibit its unique cocktail of complement regulatory proteins, we showed here that cells expressing C3 in the GBM microenvironment also upregulate several components of the classical pathway, and downregulate early complement inhibitors. These data could suggest early complement activation in areas of hypoxia, where cells expressing C3 may still be protected from detrimental effects of complement activation through upregulation of the late complement inhibitor CD59.

Therapeutic targeting of the complement system has shown to be somewhat challenging, due to the risks associated with tissue damage or infection when complement is dysregulated [56]. C3a/C3aR-signaling has been shown to be of importance in general brain physiology, including neural development [57], response to hypoxic-ischemic injuries [58], and as an important regulator of blood brain barrier integrity and the brain vascular network [59, 60]. Importantly, C3a/C3aR signaling has been associated with stemness and properties such as migration, proliferation, and angiogenesis in several cancers [61]. Due to its high expression in gliomas, we hypothesized and demonstrated that the C3a/C3aR-signaling axis could be a suitable treatment target in GBM. Our findings that radiotherapy combined with C3aR inhibition could extend the lifespan of glioma-bearing mice are in line with recent work demonstrating efficacy of combining radiotherapy with C3a inhibition in preclinical models of pancreatic cancer [62].

Previous studies have shown that tumor associated macrophages (TAMs) are a major population of the TME in gliomas, and are key contributors to several aspects of glioma progression. Attempts to deplete or re-educate TAMs through CSF-1/CSF-1R inhibition have shown promising results in pre-clinical studies, and are currently in clinical development [63]. Pyonteck et al. showed that inhibition of CSF-1R resulted in a blockage of glioma progression by reducing the number of M2-polarized macrophages, while preserving the total amount of TAMs [64]. Recent work highlighted a subset of F4/80high/CD115high/C3aRhigh/CD88high TAMs that are recruited to melanoma tumors through the NF-KB1-CSF1R-C3aR axis [65]. Anti-CSF1R therapy resulted in reduced tumor volume, F4/80 infiltration, and proliferation of macrophages, as well as blood vessel formation [65]. Similarly, blocking of a subset of TAMs positive for *C3AR1*, *CXCR4* and *CSFR1*, using either anti-C3a therapy or SB290157 treatment of B16 melanoma mice, resulted in reduced F4/80 infiltration as well as slowed tumor growth [66]. In our glioma model, SB290157-treated tumors had fewer M2-polarized macrophages as measured by CD206 expression, but no general reduction in macrophage/microglia content, as measured by F4/80. CD206+ macrophages were associated with perivascular niches of the tumors both with and without irradiation. SB290157-treated tumors displayed a trend towards lower proliferation of F4/80+ cells, and tumors treated with SB290157 in combination with radiotherapy were less vascular as measured by CD34 immunostaining. These data are in line with previous reports suggesting that therapies targeting CSF-1/CSF1R and C3a/C3aR result in altered TAM polarization, recruitment, proliferation, and altered vascularity through regulation of VEGF. Previous reports established that C3aR-deficient or SB290157-treated mice display reduced vascular density [67, 68], and that C3aR+ TAMs located in perivascular niches display an M2-like character, and are associated with regulation of angiogenesis and tumor aggressiveness [65, 66, 69, 70].

Taken together, our findings suggest that C3a/C3aR could be a viable therapeutic target in GBM, targeting of which could affect both tumor cells per se, as well as the immunosuppressive TME of GBM.

## Author contributions

RR, KIS, JS, AMB, and AP designed the study

RR, KIS, EJ, CB, JW, VP, and CC conducted experiments

RR, and JS acquired data

RR, KIS, JS, CB, JW, AMB, and AP analyzed data

CC, KP, AMB, and AP supervised the work

RR and AP drafted the manuscript

All authors reviewed, edited, and approved the final version of the manuscript

## Acknowledgements

Funded by the European Union (ERC, RESISTANCEPROGRAMS, 101043587). Views and opinions expressed are however those of the author(s) only and do not necessarily reflect those of the European Union or the European Research Council Executive Agency. Neither the European Union nor the granting authority can be held responsible for them. This study was further supported by the Ragnar Söderberg Foundation, the Swedish Cancer Society, the Swedish Research Council, the Swedish Childhood Cancer Fund, Ollie & Elof Ericssons foundation, and the Crafoord foundation.

