## Supplementary Figures and Legends for "Hypoxia-induced Complement Component 3 Promotes Aggressive Tumor Growth in the Glioblastoma Microenvironment"

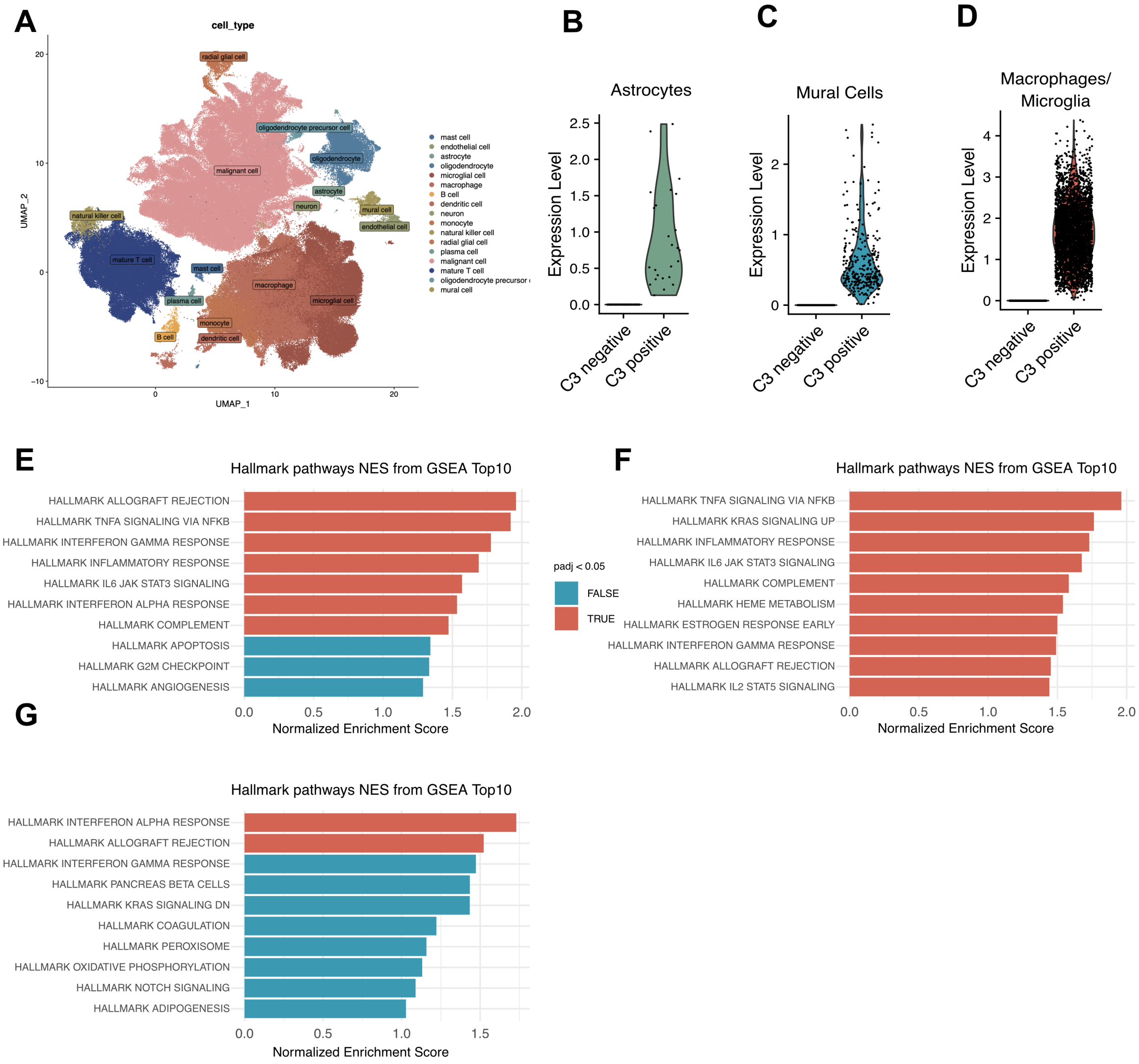


***Supplementary Figure 1.***

***A,*** *GBmap from Ruiz-Moreno, et al. 2022 (BioRxiv) comprising 16 data sets from 110 patients.* ***B-D****, C3 transcript level subdivided into C3-expressing or non-expressing cells from astrocytes, mural cells and macrophages/microglia.* ***E-G****, Hallmark enriched gene signatures in* $C3^{+}$ *cells from astrocytes, mural cells and macrophages/microglia origin. Red- and blue-colored bars indicate significant and non-significant Benjamini-Hochberg adjusted P values (padj < 0.05), respectively.*

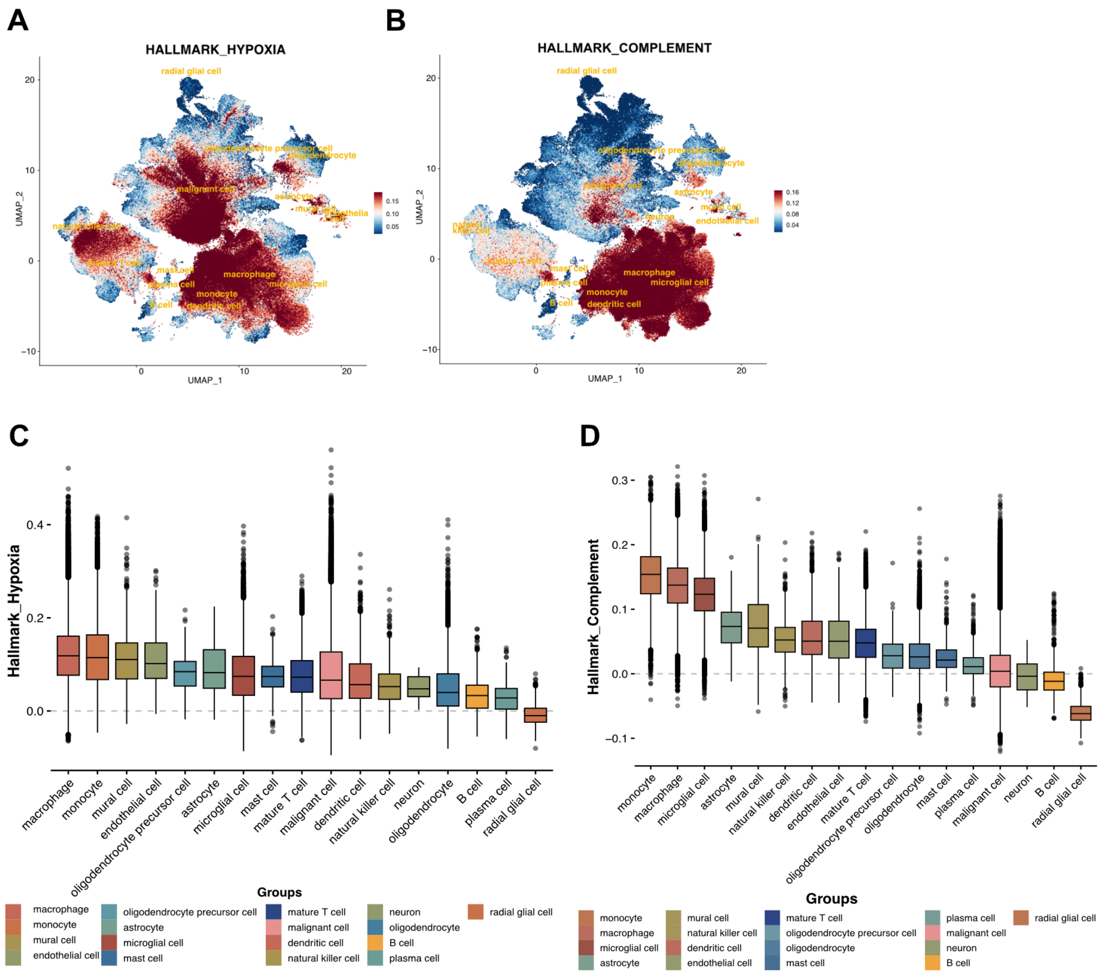


***Supplementary Figure 2.***

***A,*** *UMAP displaying Hallmark hypoxia gene signature mapped onto GBmap.* ***B,*** *UMAP displaying Hallmark complement gene signature mapped onto GBmap.* ***C-D,*** *Cell types expressing hypoxic and complement gene signatures in GBM. Color scale in A-B and values in C-D indicate hypoxia and complement module scores.*

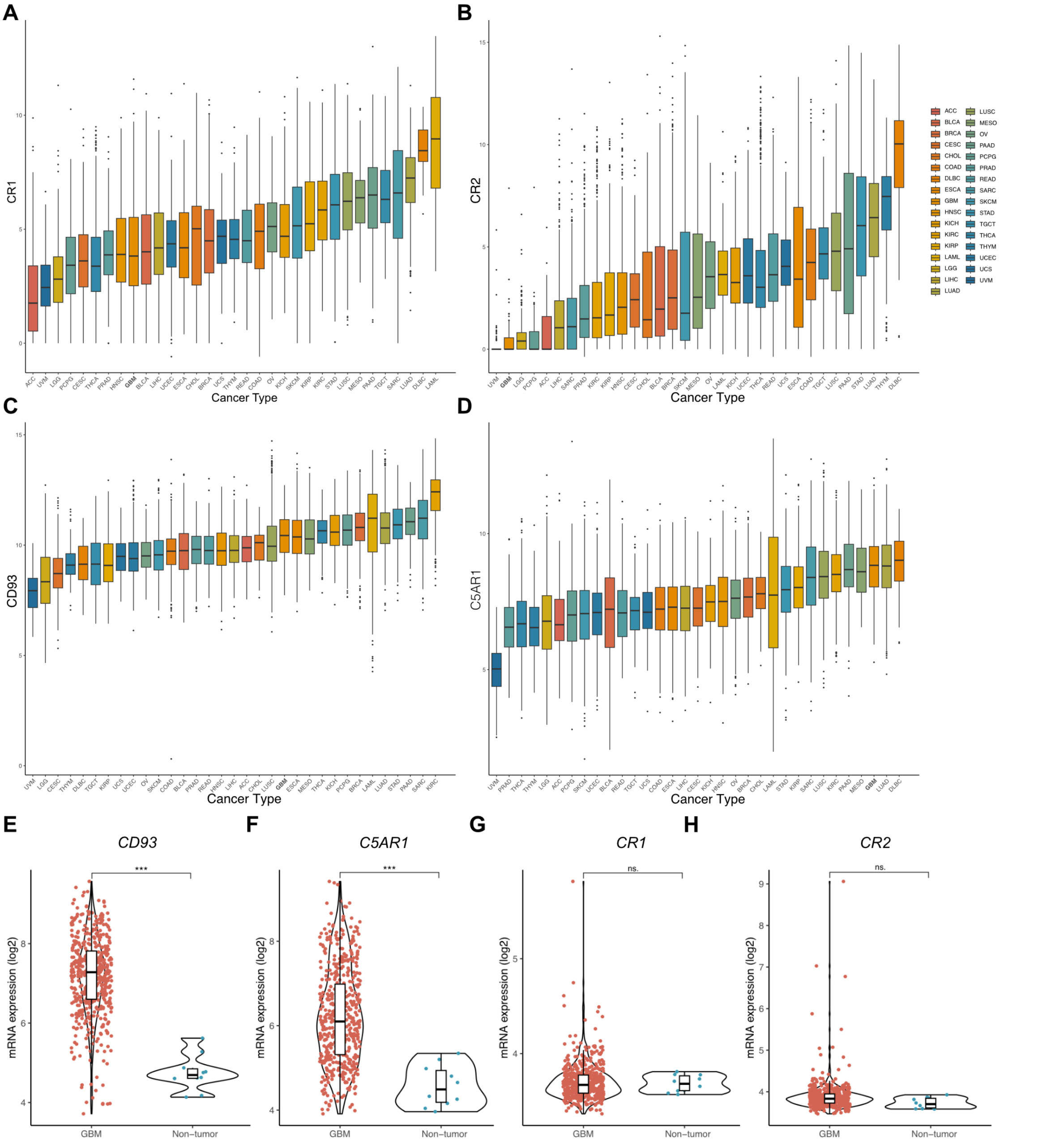


***Supplementary Figure 3.***
***A-D,*** *CR1, CR2, CD93 and C5AR1 transcript expression of Pan-Cancer TCGA data of common cancer types (n=33).* ***E-H,*** *CR1, CR2, CD93 and C5AR1 transcript expression in GBM compared to normal tissue. Statistical analysis were performed with Tukey's Honest Significant Difference (HSD).*

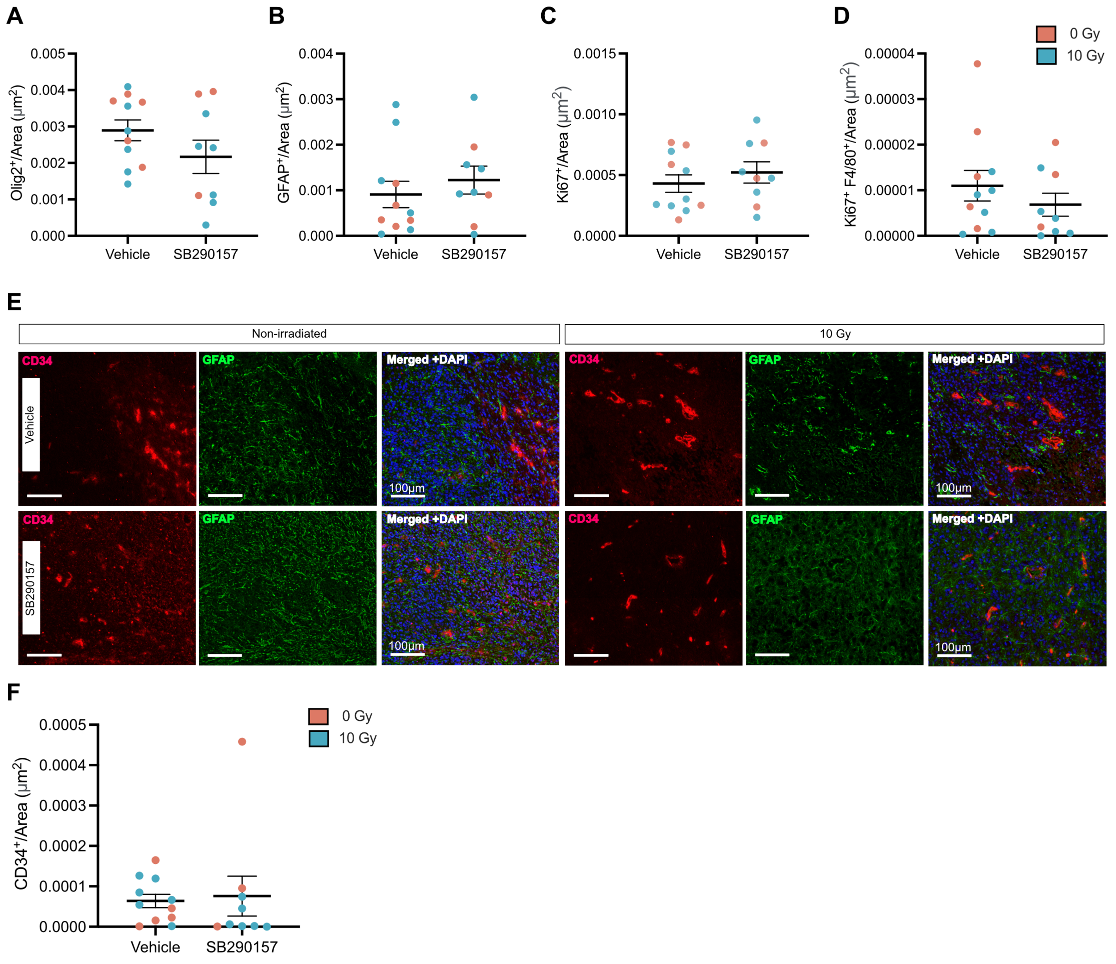


***Supplementary Figure 4.***

***A-B,*** *Quantitative analysis of $Olig2^+$ or $GFAP^+$ cells/Area ($\mu m^2$) from Vehicle or SB290157 from immunofluorescent staining in figure 5C.* ***C,*** *Representative immunofluorescent staining of vessels (CD34) and astrocytes (GFAP) in SB290157 treated mice.* ***H,*** *Quantitative analysis of $CD34^+$ cells/Area ($\mu m^2$) from Vehicle or SB290157. Statistical analysis were performed with Mann-Whitney U test for treatments. *, P < 0.05, **, P < 0.01, or ***, P< 0,001.*
